# Four Core Genotypes Mice Exhibit Quantitative Differences in T and B Cell Subpopulations compared to Wild-type Mice

**DOI:** 10.64898/2026.02.16.706181

**Authors:** Gillian R. McGuire, Abigail S. Klimas, Daniel F. Deegan, Gennaro Calendo, Catherine F. Alapatt, M. Raza R. Zaidi, Andrea Bottaro, Nora Engel

## Abstract

It has been well established that females have a more active immune system. Females respond better to vaccines, are more resistant to somatic cell cancers, display better pathogen responses, present antigens better, and, conversely, are more prone to autoimmune diseases compared to male counterparts. Though these trends have been observed across normal and pathogenic states, the mechanisms underlying these sex differences have not been fully explained. Some hormonal effects on immune cell populations have been reported, but much less is known about effects contributed by genes on the sex chromosomes, for example those that are more highly expressed in females due to X inactivation escape, or Y-linked genes those unique to males. Here we use the Four Core Genotypes (FCG) mouse model and wildtype XY male mice to disentangle the effects of sex hormones, sex chromosome complement, and their interactions on baseline B and T cell populations in the periphery and T cells in the thymus. We test the effects of a previously described X-Y chromosomal translocation and those of the *Sry* transgene insertion on chromosome 3. We observe that mice harboring the *Sry* transgene show significant depletion of peripheral CD8+ T cell subpopulations. In the thymus, the XY XY,but not the XX males, show significant decrease to both CD8+ and CD4+ single positive T cells and an increase to CD4/CD8 double positive T cells. We also show that Y chromosome-bearing mice exhibit depletion in splenic marginal zone B cells. Our data suggests that the gonadal sex is the strongest contributor to this phenotype. Our studies define a critical framework for the use of this model and provide valuable data to assess the use of the FCGs model, especially for diseases involving the immune response.

## Introduction

Sex is defined as a biological variable based on the organization of sex chromosomes, the development of reproductive organs, and the level of sex steroid hormones. The complex interplay between these factors can contribute to sex biases in any phenotype. For example, it has been well established that there are prominent sex differences in the immune system^1–3^. Overall, females tend to have more active immune responses than males^4^, resulting in better protection against infectious pathogens and increased vaccine efficacy^5^, but also higher rates of autoimmune disease^4,6,7^. In pathogen response, it is known that females detect pathogens due to higher expression of pathogen-associated molecular pattern (PAMP) receptors, along with better phagocytosis, direct bacterial killing, and antigen presentation from antigen-presenting cells (APCs)^8–10^. Similar trends and mechanisms have been observed in anti-tumoral responses in reproductive and non-reproductive cancers, where for example, androgen receptor blockade in prostate cancer robustly increased the abundance and anti-tumor functionality of tumor infiltrating T cells^11,12^. Estrogen itself has a paradoxical role in anti-tumor responses, where it can be immune suppressive and immune activating, dependent on the receptors and evasion mechanisms upregulated by cancer cells^12,13^. In non-small cell lung cancer, estrogens contribute to immune evasion. In contrast, they exert anti-tumoral signaling by inducing the release of IFN-γ^14^. Females’ immune systems are also generally thought be better primed against cancer due to strong immune activation mechanisms with antigen presentation, T cell activation, and boosted effector cell functionality^12^. However, the mechanisms underlying these differences are still unknown.

Sex hormones are one contributing factor for the sex differences in the immune system. Both androgens and estrogens are major regulators of immune responses, and their receptors are expressed in both innate and adaptive immune system cells^15^. Estrogen, circulating as 17β-Oestradiol (E2), can bind to three receptors which are: ERα, ERβ, and GPER1, all of which are expressed on immune cells in varying quantities^4^. Intriguingly, male and female T and B lymphocytes have the same levels of ERα and ERβ, underscoring that the levels of circulating hormones may be critical in regulating immune cell function^4,16^. In T cells, ERα signaling is important for differentiation into T_reg_ lineages and in mature T cell cytokine secretion^15^. In the context of anti-tumor immunity, it is known that female dendritic cells are better at presenting antigens to T cells than in males, which may promote better immune infiltration, activation, and thus killing of cancer^17,18^. However, how this influences T cell relevant pathologies such as autoimmune diseases and cancer remains unclear.

Androgens, which circulate as testosterone or DHT, are overall thought to be immune suppressive^5^. Androgens play a role in the NF-κB pathway by increasing expression of PPARα in T cells to suppress inflammatory activation^19^. Lack of testosterone in mice, such as following castration, alters T cell subpopulations by increasing CD4+ and CD8+ numbers, which highlights the role of an androgen microenvironment in suppressing T cell immunity^20^. Overall, the field has made advancements into understanding how the sex steroid hormones play a role in baseline immunity, but further research is essential to bridge this to various immune implicated pathologies.

Another less explored factor contributing to sex differences in immunity involves proteins encoded on the X and Y chromosomes. Some genes relevant to the immune system escape X chromosome inactivation, causing dosage differences between females and males, while expression of Y chromosome-specific genes is restricted to males^21^. The importance of X chromosome genes in immune function is highlighted by the large number of X-linked primary immunodeficiency diseases^21^, but also by sex-related immune effects caused by gene dosage differences. For example, the DDX3X gene that escapes X inactivation is critical in IFN-γ production pathways which combat certain pathogens such as *Listeria monocytogenes*^*22,23*^. The IFN-γ response is also critical in anti-tumoral immunity^24^. In immunogenic cancers, such as melanoma, it has been linked to improved survival in females^25^. Loss of function mutations in melanoma in males has also been linked to transcriptional signatures associated with invasiveness and metastasis, events that are often dependent on immune evasion^26,27^.

A less extensively studied source of sex differences relates to the function of Y chromosome encoded genes. It has been shown that Y-linked genes are critical in immune gene regulation in mice^28^. In the context of anti-cancer immunity, the loss of Y chromosome is associated with a CD4+ Treg phenotype and a loss of cytotoxic effector function in CD8+ T cells^29^. Overall, loss of Y in infiltrating CD4+ and CD8+ T cells may promote immunosuppression and reduced cancer killing^30^.

The Four Core Genotypes (FCG) mouse model was developed as a tool to disentangle whether sex biases observed in a specific phenotype could be attributed to the effect of the sex chromosomes, of gonadal sex hormones, or both^31^. *Sry* is the single fate-determining gene in male gonadal sex development during embryonic development^32^. The FCG founder male combines *Sry* as a transgene on chromosome 3 with the deletion of the Y chromosome-linked *Sry* gene, to separate the Y chromosome from male gonadogenesis^31,33^. When this founder male, designated XY-*Sry*+, is bred with a wildtype C57BL/6 XX female, four genotypes of mice can be produced which are the following: XX females, XY^−^ females, XX*Sry*+ males, and XY^−^ *Sry*+ males^31^. This model has been used to address a variety of sex-biased conditions, such as cardiovascular disease and neurological development, but has only rarely been leveraged to study cancer and immune oncology^31^. For example, in bladder cancer the X-linked *Kdm6a* was implicated in a protective epigenetic mechanism in females^34^. Thus, the FCG model has potential to uncover independent mechanisms one cannot see in wildtype male versus female comparisons in physiology and pathology.

There are however some genetic features of the FCG model that raise issues for the mechanistic interpretation of findings in this model. First, the *Sry* transgene consists of multiple copies, not all of which are functional, integrated at Chr3 70673749-70673824 bp (NCBI37/mm9)^33^. A 74bp deletion of chromosome 3 is present at the integration site, which does not interrupt known protein coding genes or pseudogenes^31^. Expression of the genes neighboring the *Sry* transgene in mouse liver do not appear to be significantly affected, but two genes on chromosome 3 (*Ppid* and *Lxn)* were found to be altered^33^. The gene *Ppid* is approximately 9 million bp upstream from the insertion of *Sry*, and *Lxn* is about 3 million bp downstream from the insertion (NCBI37/mm9). The effect of transgenic *Sry* in other adult tissues, including immune cells, is not known. We also do not know if expression of neighboring genes on chromosome 3 is affected in other tissues.

Furthermore, it was recently reported that the Y chromosome lacking the *Sry* gene in the XY^**-**^ *Sry*^**+**^ founder male contains nine genes translocated from the X chromosome, thus altering their dosage in Y-chromosome-bearing mice compared to wild-type XY males^35^. It is not known if and in which tissues these genes are expressed and lead to a phenotypic consequence, but some of them, such as *Tlr7* and *Tlr8*, are immune-related and known to escape X inactivation in some immune cell subpopulations in humans^36,37^. The effect of these unique genetic alterations in FCG mice on immune system parameters and function is unknown but must be considered when evaluating immune phenotypes in this model^38^.

Here we report significant alterations in peripheral immune subsets in FCG mice that appear linked to the presence, and possibly dysregulated expression, of the FCG *Sry* transgene, or due to the X to Y chromosome translocation in these mice.

## Methods

### Mouse Colony

Mouse studies were performed under approval of Cooper University Health Care Institutional Animal Care and Use Committee. Mice were maintained under standard conditions of 12-h light and dark cycles (22°C ± 1°C, with food and water *ad libitum*). We used the Four Core Genotypes (FCG) mouse model^31,39,40^ on a C57BL/6J background (B6.Cg-Tg(Sry)2Ei *Sry*^*dl1Rlb*^ T(XTmsb4x-Hccs;Y)1Dto/ArnoJ, Jackson Laboratories stock 010905; backcross generation greater than 20) (*a generous gift of Arthur Arnold*). The founder males in this model have the *Sry* gene deleted from the Y chromosome and inserted as a transgene on chromosome 3 (XY^**-**^ *Sry*^**+**^, referred to here as FCG XY males). Crossing FCG XY males with wildtype XX females results in XX and XY mice with ovaries and XX and XY mice with testes, dependent solely on whether the *Sry* transgene is inherited and independently of the presence of the Y chromosome (**Figure 1**). Thus, the gonadal type of the progeny (testes or ovaries) is independent of the sex chromosome constitution. This strain also contains 9 genes translocated from the X chromosome to the Y (*Tmsb4x, Tlr8, Tlr7, Prps2, Frmpd4, Msl3, Arhgap6, Amelx* and *Hccs*). Analyzing these four core genotypes allows distinguishing between sex chromosome and sex hormone dependent effects. We also utilized C57BL/6J wildtype XY males (Jackson Labs, Stock #000664) for comparison to the FCG line. We will refer to the C57BL/6J XY males as wildtype XY males.

**Figure 1:**
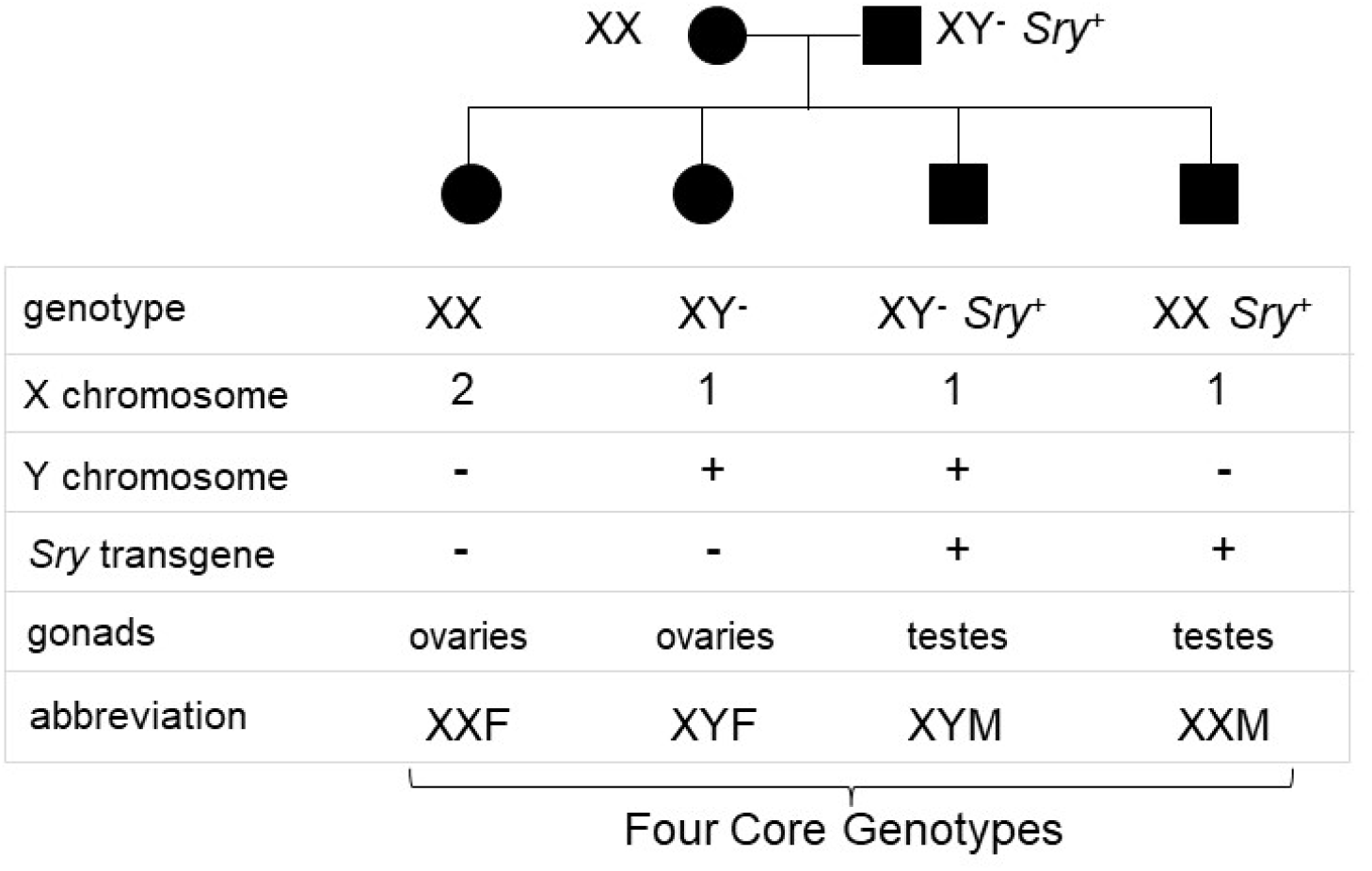
The Four Core Genotypes (FCG) mouse model decouples the Y chromosome from male gonadogenesis.

8-15 week old mice were used for studies. Necropsy was performed to extract axillary and inguinal lymph nodes bilaterally, spleen, and thymus. Fresh tissue was placed in a cell culture medium (DMEM with 10% FBS, 1% L-glutamine, and 1% 100X antibiotic/antimycotic solution) or FACS buffer (1X PBS, 1-5% FBS, depending on the organ) on ice and immediately processed for flow cytometry.

Using 8-12 week old mice, approximately 100-200uL (dependent on mouse weight), of whole tail vein blood was collected in RAM Scientific SAFE-T-FILL Micro Capillary Blood Collection: EDTA K2 tubes (Fisher Scientific: #14-915-50). Samples were placed on ice and shipped to Fox Chase Cancer Center (Dr. Kerry Campbell) within 4 hours of collection for flow cytometry processing.

### FCG Genotyping

DNA was extracted from ear punches from adult mice and genotype was determined by PCR using primers for *Sry* and *Ymt*. PCR products were then run on a 2% agarose gel for analysis. Gonadal males exhibit a positive result for *Sry*, and positive or negative for *Ymt* reveals the presence or absence of a Y chromosome. PCRs were performed using DreamTaq Green PCR Mastermix (2x) (Thermofisher #K1081). Primer sequence, annealing temperature, and product size are listed in **Table 1**.

**Table 1:**
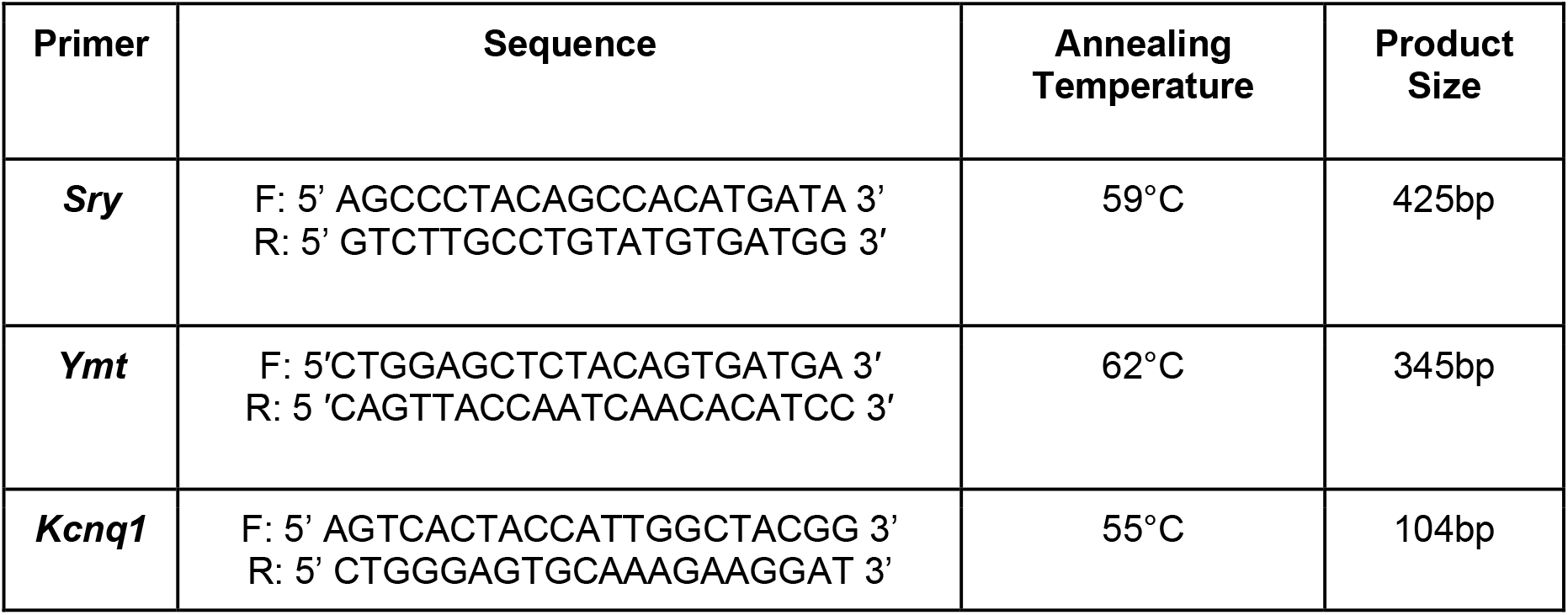
Primers for mouse genotyping.

### Flow Cytometry

#### Analysis of spleen, thymus and superficial lymph nodes (LN)

Single cell suspensions were obtained from mouse spleen, thymus and pooled axillary and inguinal lymph nodes by gentle disaggregation between frosted microscope slides in FACS buffer (1X PBS, 5% FBS). Erythrocytes were eliminated from spleen suspensions using Red Blood Cell Lysis Buffer (Abcam). Cell suspensions were filtered through 100um nylon mesh, washed twice in FACS buffer and counted. A range of 0.5-2×10^6^ cells/sample were used for flow cytometry staining using combinations of the antibodies listed below. Dead cells were excluded using LIVE/DEAD fixable Aqua Dead Cell Stain Kit (Invitrogen). Data were collected on a Stratedigm S1300 apparatus and analyzed using CellCapture software. The antibodies used are in **Table 2**. The analysis setup is **Figure 2**.

**Table 2:** Antibody panels for flow cytometry analyses.

| Biomarker | Fluorophore | Source (catalog number) | μl/sample |
| --- | --- | --- | --- |
| CD8 | FITC | BioLegend<br>#100803 | 2 |
| NK-1.1 | PE | BioLegend<br>#156503 | 2 |
| CD4 | Pacific Blue | BioLegend<br>#100534 | 3 |
| CD45 | BUV395 | BD #564279 | 1 |
| FVS440 | AF350 | BD #566332 | 1 (of 1/25 dilution) |
| CD3 | BV711 | BioLegend<br>#100241 | 4 |
| CD19 | FITC | BD #557398 | 0.5 |
| CD21/35 | Pacific Blue | Biolegend #123414 | 1 |
| CD23 | PE-Cy7 | BioLegend #1011614 | 1 |
| CD4 | FITC | BD #553729 | 0.5 |
| CD3 | PerCP-Cy5.5 | BD #551163 | 1 |

Table 2: Antibody panels for flow cytometry analyses
| Biomarker | Fluorophore | Source (catalog number) | µl/sample |
| --- | --- | --- | --- |
| CD8 | FITC | BioLegend<br>#100803 | 2 |
| CD8 | AF647 | BD #557682 | 1 |
| CD45 | APC-Cy7 | BioLegend #103116 | 1 |

**Figure 2:**
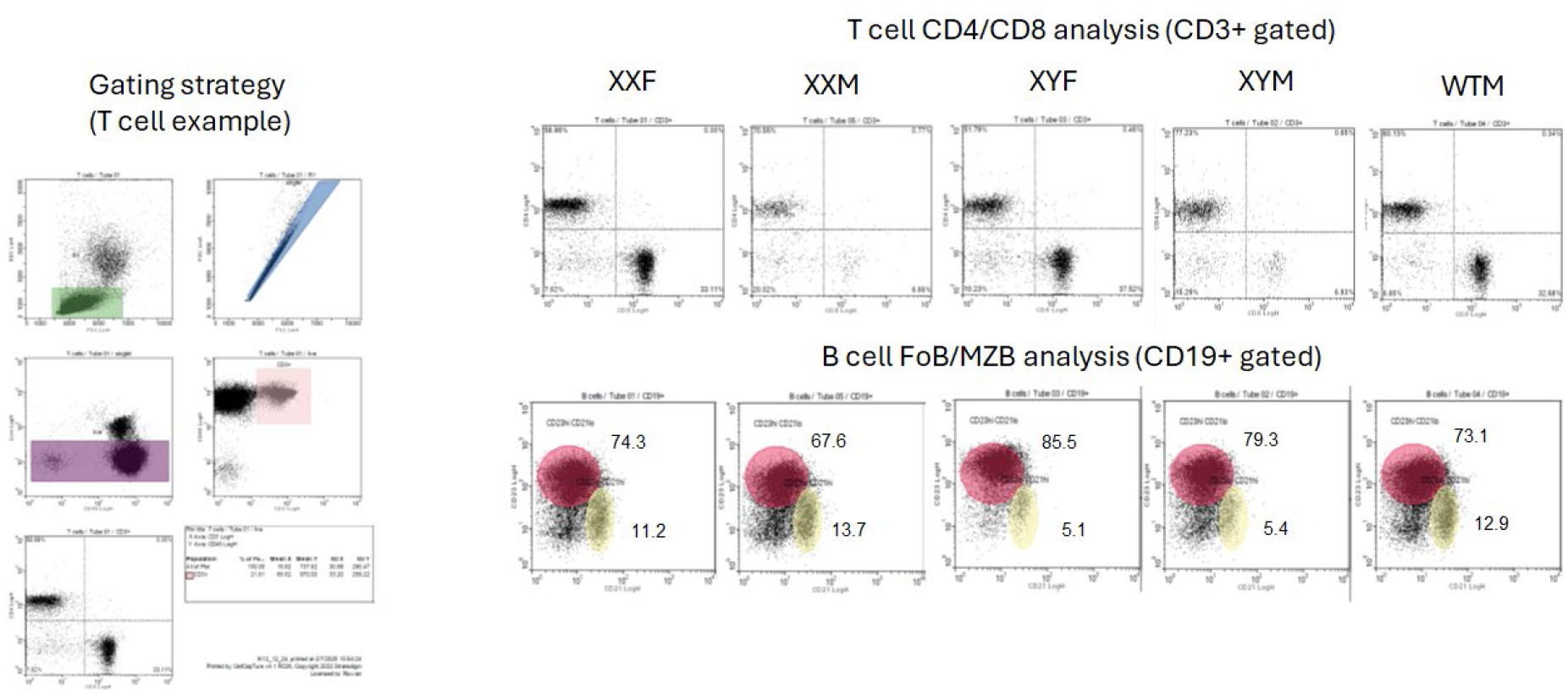
Flow Cytometry gating strategy for B & T Cells in Four Core Genotypes (FCG) & wildtype C57BL/6J murine tissues.

#### Analysis of whole tail vein blood

Whole tail-vein blood (50-200 µl) was mixed with 3 ml ice cold erythrocyte lysis buffer (0.15 M ammonium chloride, 10 mM potassium bicarbonate, and 0.1 mM disodium EDTA) and incubated on ice for 5 minutes. Lysis was terminated by immediately adding 12 mL of ice cold RPMI-1640 deficient medium (lacking FBS, biotin, riboflavin, and phenol red) and centrifuging at 500xg for 5 minutes at 4°C. The cell pellet was resuspended in 5 ml of ice cold RPMI-1640 deficient medium, centrifuged again, and resuspended in 100-200 µl of RPMI-1640 deficient medium. Viability was determined with trypan blue dye and mixed with antibodies in staining tubes, gently vortexed, and incubated on ice for 20 minutes. Cells were washed on ice once with 3 ml of FACS wash buffer (1% heat inactivated FBS and 0.9% sodium azide in Hanks’ Balanced Salt Solution), once with 3 ml of wash buffer containing propidium iodide (20 µl of 1 mg/ml stock/500 ml), and resuspended in 100-200 µl of wash buffer before analysis on a BD FACSymphony S6 spectral flow cytometer in the Fox Chase Cancer Center Cell Sorting Facility. Data was analyzed with FlowJo (version 10.10.0). The antibodies used are in **Table 2**. The analysis setup is in **Figure 3**.

**Figure 3:**
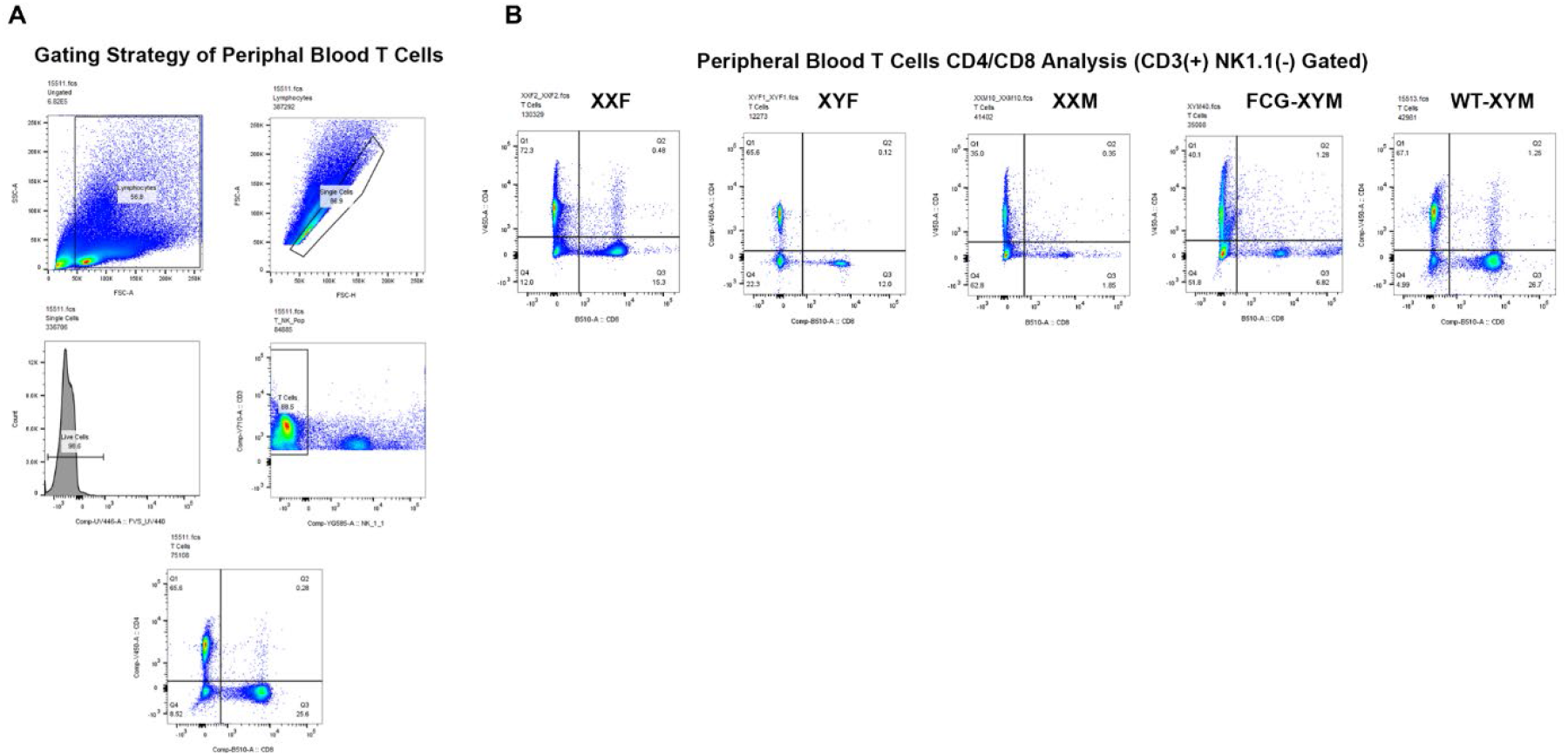
Flow cytometry gating strategy for T cells in Four Core Genotypes (FCG) & wildtype C57BL/6J peripheral blood. (A) General gating strategy for peripheral blood T lymphocytes (wildtype C57BL/6J male used as an example), and (B) Peripheral blood C

## Results

### FCG Males Have a Peripheral Reduction of CD8+ T Lymphocytes

To first assess immunophenotype differences that are relevant to anti-tumoral immunity in Four Core Genotypes (FCG) mice, we investigated T lymphocytes in peripheral blood, spleen, superficial lymph nodes (axial and inguinal), and thymus. In most rodent species, including *Mus musculus*, peripheral blood contains approximately 70–80% lymphocytes^41^. We noticed a significantly lower fraction of T lymphocytes (CD45+ CD3+) in FCG males, regardless of host sex chromosome type (XX or XY), in comparison to wildtype XY males, wildtype XX females, and FCG XY females (**Figure 4**). Compared to wildtype XY males, FCG XY males have a 50-60% decrease and FCG XX males a 60% decrease in their overall T lymphocytes (**Figure 4-A**). This decrease primarily affects the CD8+ T lymphocyte subpopulation (**Figure 4-B & C**), which shows an 80-90% reduction compared to wildtype XY male counterparts (**Figure 4-B**).

**Figure 4:**
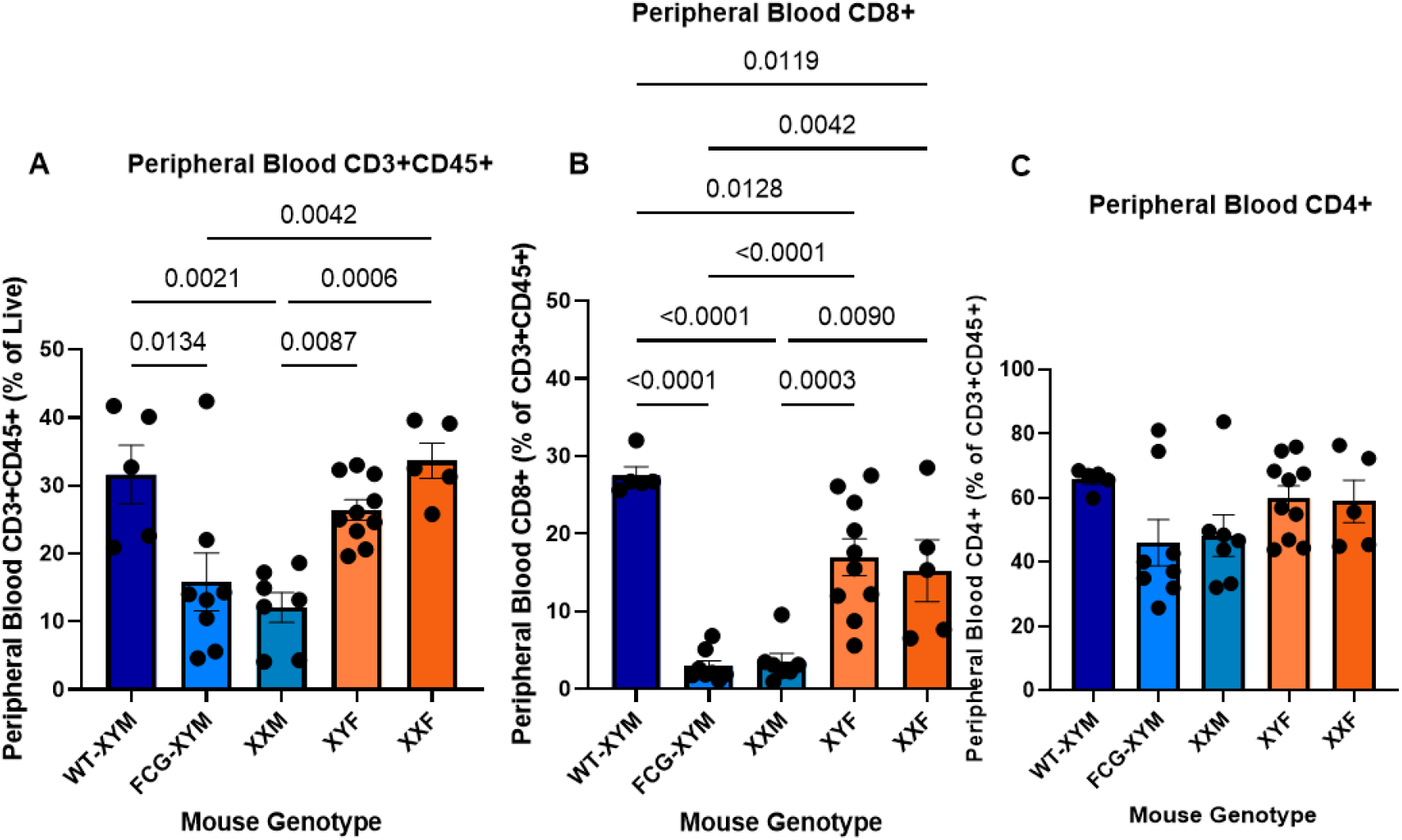
T cell subpopulations in peripheral blood between Four Core Genotypes (FCG) mice and wildtype (WT) C57BL/6J males. (A) Peripheral blood CD3+CD45+ total T lymphocytes (% of live). (B) peripheral blood CD8+ T lymphocytes (% of CD3+CD45+), and (C) peripheral blood CD4+ T lymphocytes (% of CD3+CD45+). For (A) to (C): N_WTM_=5, N_XYM_=8, N_XXM_=7, N_XYF_=10, N_XXF_=5. Male gonadal sex marked in blue; female gonadal sex marked in orange. All stats One-way ANOVA plus Tukey-HSD on the means (significance at p<0.05), error bars SEM. Significance bar not shown defines p≥0.05, which is nonsignificant (NS).

To expand these findings, we looked at the immune cell composition of lymphoid tissues, including thymus and peripheral lymphoid organs (spleen and superficial - axillary and inguinal - lymph nodes). Results in the lymph nodes and spleen recapitulate the results in peripheral blood (**Figure 5**). In the spleen, FCG males have a 70-80% decrease in their overall T lymphocyte (CD3+) populations and a 50-70% decrease in the CD8+ T lymphocyte cluster compared to their wildtype XY male counterparts (**Figure 5-A & B**). In the superficial lymph nodes (LN), FCG males exhibit a 50-60% decrease in their overall T lymphocyte (CD3+) populations and a 60-70% decrease in the CD8+ T lymphocyte cluster compared to their wildtype XY male counterparts (**Figure 5-D & E**). In both cases, there are no differences in the CD4+ T cell subpopulation percentages (**Figure 5-C & F**). FCG XY females and wildtype XX females only show differences to wildtype XY males in the peripheral blood CD8+ T cell population (**Figure 5-B & E**) (**Figure 3-B**). Wildtype XX females and XY females have a 40% reduction in CD8+ T cells in the circulating blood compared to wildtype XY males (**Figure 5-B & E**) (**Figure 3-B**).

**Figure 5:**
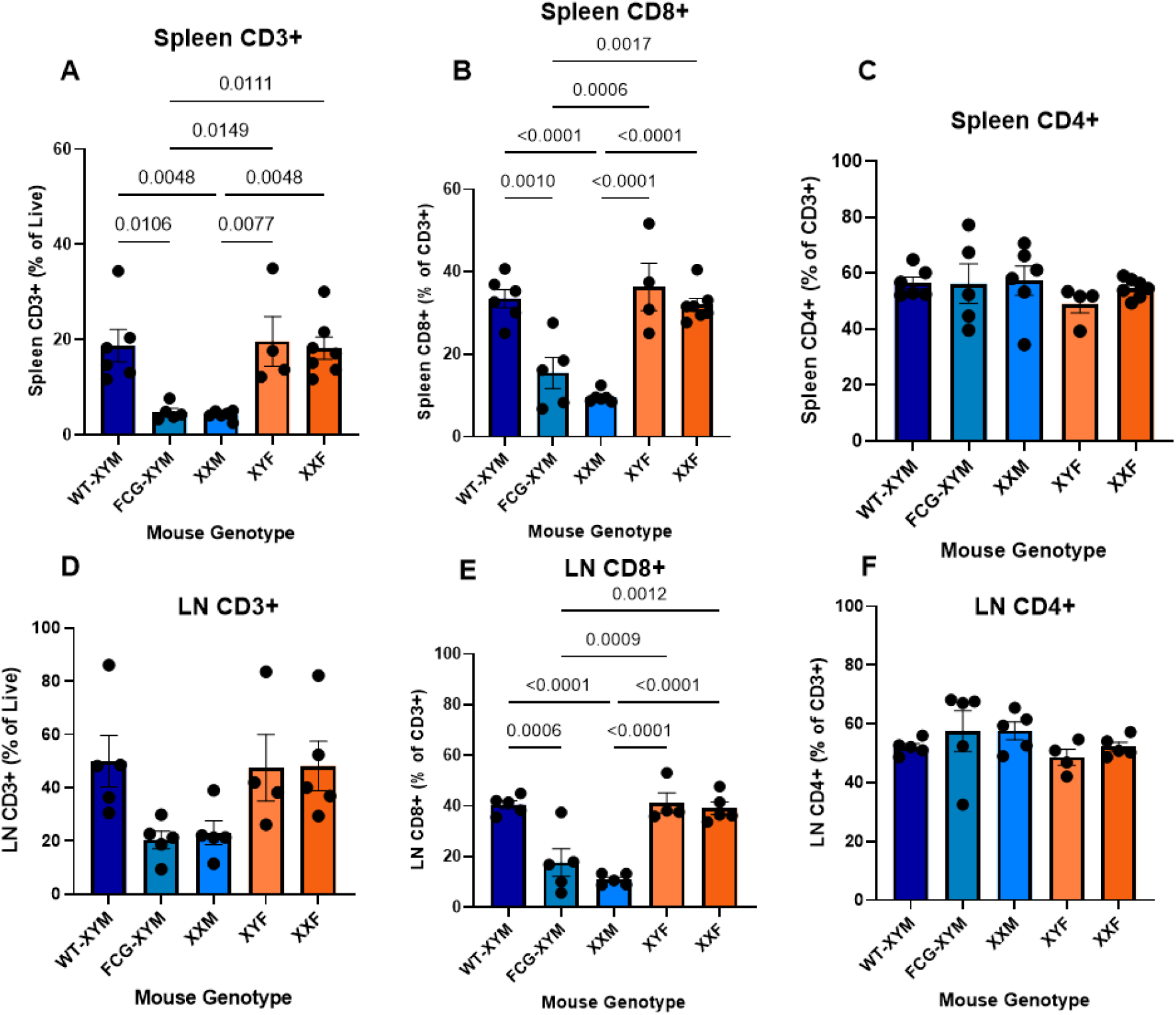
T cell subpopulations in spleen & superficial lymph nodes (LN) between Four Core Genotypes (FCG) mice and wildtype (WT) C57BL/6J males. For (A)-(C), analysis of spleen. For (D)-(F), analysis of LN. (A) spleen CD3+ total T lymphocytes (% of live), (B) spleen CD8+ T lymphocytes (% of CD3+), (C) spleen CD4+ T lymphocytes (% of CD3+), (D) LN CD3+ total T lymphocytes (% of live), (E) LN CD8+ T lymphocytes (% of CD3+), and (F) LN CD4+ T lymphocytes (% of CD3+). For (A) to (C): N_WTM_=6, N_XYM_=5, N_XXM_=6, N_XYF_=4, N_XXF_=7. For (D) to (F): N_WTM_=5, N_XYM_=5, N_XXM_=4, N_XYF_=4, N_XXF_=5. Male gonadal sex marked in blue; female gonadal sex marked in orange. All stats One-way ANOVA plus Tukey-HSD on the means (significance at p<0.05), error bars SEM. Significance bar not shown defines p≥0.05, which is nonsignificant (NS).

To investigate whether these peripheral T cell subset differences are due to maturation defects, we investigated the composition of the thymus (**Figure 7**). Unexpectedly, only the FCG XY males show significant differences to T cells in the thymus (**Figure 7**). FCG XY males show significant decrease to CD3+ cells overall, due to decrease in the CD8+ and CD4+ single positive subpopulations, in FCG XY males compared to FCG XX males, FCG XY females, and wildtype (XY male and XX female) counterparts (**Figure 7-A & C**). We also see a slight but significant increase to the FCG XY males in CD4/CD8 double positive (DP) cells compared to all genotypes (**Figure 7-B & D**). We conclude that the majority of the CD8+ differences are restricted to the periphery, though there is a unique difference in the FCG XY male thymus (**Figure 4**) (**Figure 5**) (**Figure 7**).

One unexpected finding was the effect on peripheral double negative (DN) T cells, an overall rare peripheral population in humans and mice whose role in physiology and pathological states is poorly understood^42^ (**Figure 6**). FCG males show a strong increase to this subpopulation compared to wildtype counterparts and XY females in both blood and spleen (**Figure 6**). There is a 90% increase in this rare subpopulation in FCG males in peripheral blood (**Figure 6-A**) and a 70% increase in spleen (**Figure 6-B**), when compared to wildtype males.

**Figure 6:**
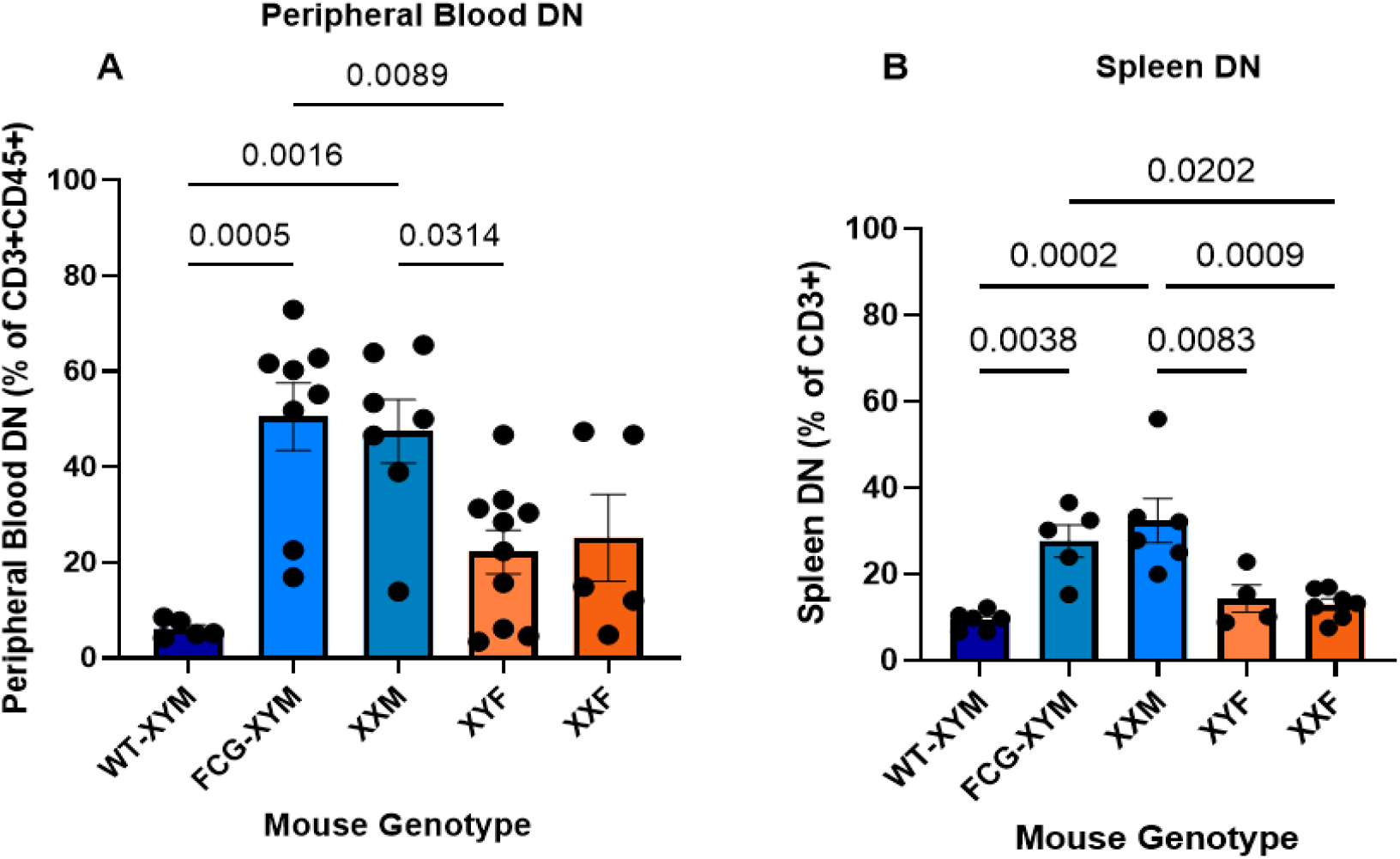
Double negative (DN) T cell subpopulations in spleen & peripheral blood between Four Core Genotypes (FCG) mice and wildtype (WT) C57BL/6J males. (A) DN T lymphocytes circulating in peripheral blood (% of CD3+CD45+), and (B) DN T lymphocytes in spleen (% of CD3+). For (A): N_WTM_=5, N_XYM_=8, N_XXM_=7, N_XYF_=10, N_XXF_=5. For (B): N_WTM_=6, N_XYM_=5, N_XXM_=6, N_XYF_=4, N_XXF_=7. Male gonadal sex marked in blue; female gonadal sex marked in orange. All stats One-way ANOVA plus Tukey-HSD on the means (significance at p<0.05), error bars SEM. Significance bar not shown defines p≥0.05, which is nonsignificant (NS).

**Figure 7:**
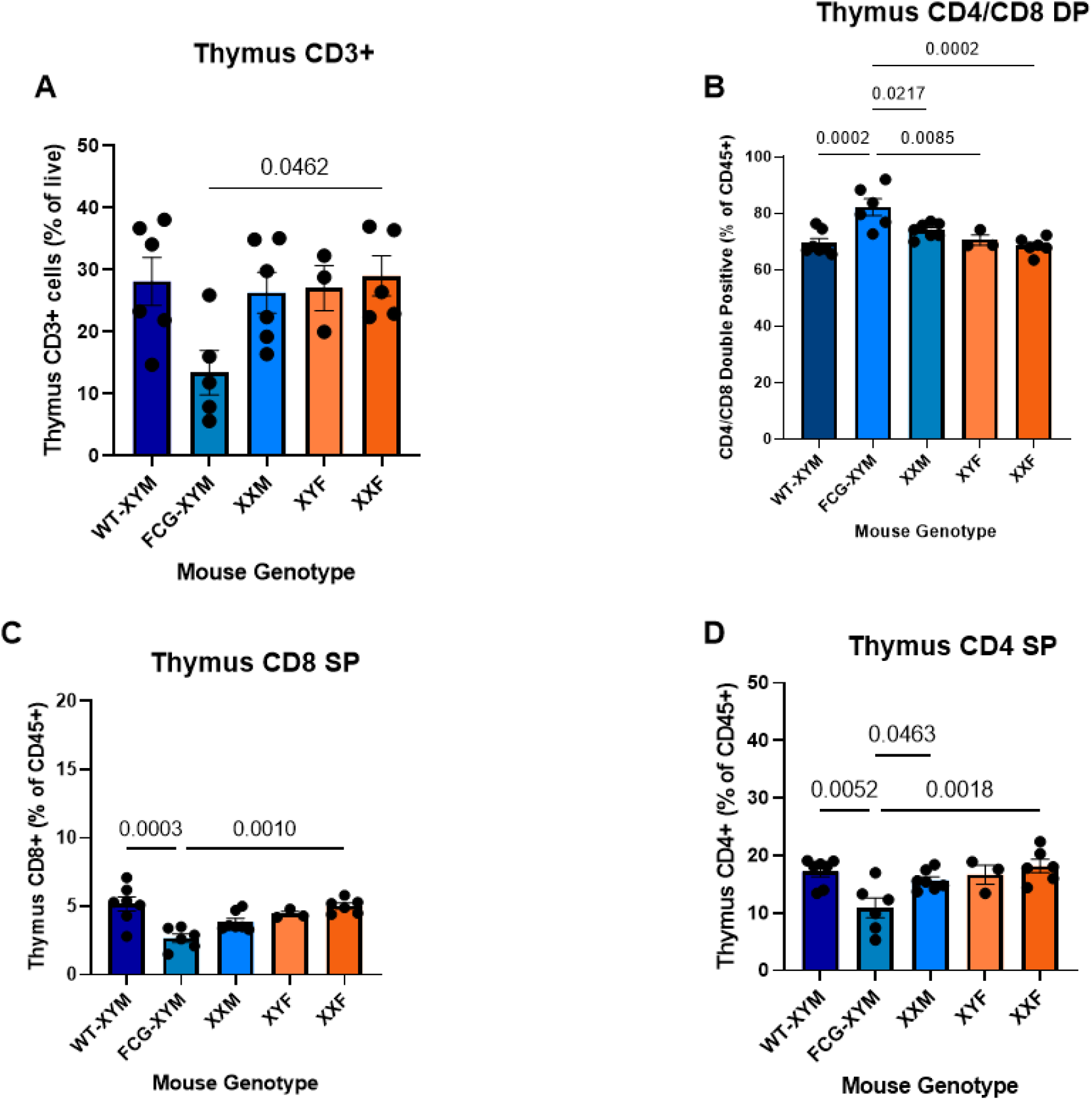
T cell subpopulations in thymus between Four Core Genotypes (FCG) mice and wildtype (WT) C57BL/6J males. (A) thymus CD3+ total T lymphocytes (% of live), thymus CD4/CD8 double positives (% of CD45+), (C) thymus CD8+ T lymphocytes (% of CD3+), and (D) thymus CD4+ T lymphocytes (% of CD3+). For (A): N_WTM_=6, N_XYM_=5, N_XXM_=6, N_XYF_=3, N_XXF_=5. For (B): N_WTM_=7, N_XYM_=6, N_XXM_=7, N_XYF_=3, N_XXF_=6. For (C): N_WTM_=7, N_XYM_=6, N_XXM_=7, N_XYF_=3, N_XXF_=6. For (D): N_WTM_=7, N_XYM_=6, N_XXM_=7, N_XYF_=3, N_XXF_=6. Male gonadal sex marked in blue; female gonadal sex marked in orange. All stats One-way ANOVA plus Tukey-HSD on the means (significance at p<0.05), error bars SEM. Significance bar not shown defines p≥0.05, which is nonsignificant (NS).

### FCG Mice Bearing the Y Chromosome Show a Significant Increase in the Ratio of Marginal Zone (MZ) B Cells and Follicular Zone (FoZ) B Cells

parallel to the analysis of splenic T cells, we also characterized B cell populations in the spleen and LN (**Figure 8**). In mice harboring the FCG Y chromosome (with the X-derived 9-gene translocation), that is FCG XY females and XY males, MZ B cells are significantly decreased by 50% compared to wildtype males, and XX bearing mice (**Figure 8-A**). This corresponds to a 2.5-fold increase to the FoZ/MZ B cell ratio in the spleen compared to wildtype XY males, and all XX mice (**Figure 8-B**).

**Figure 8:**
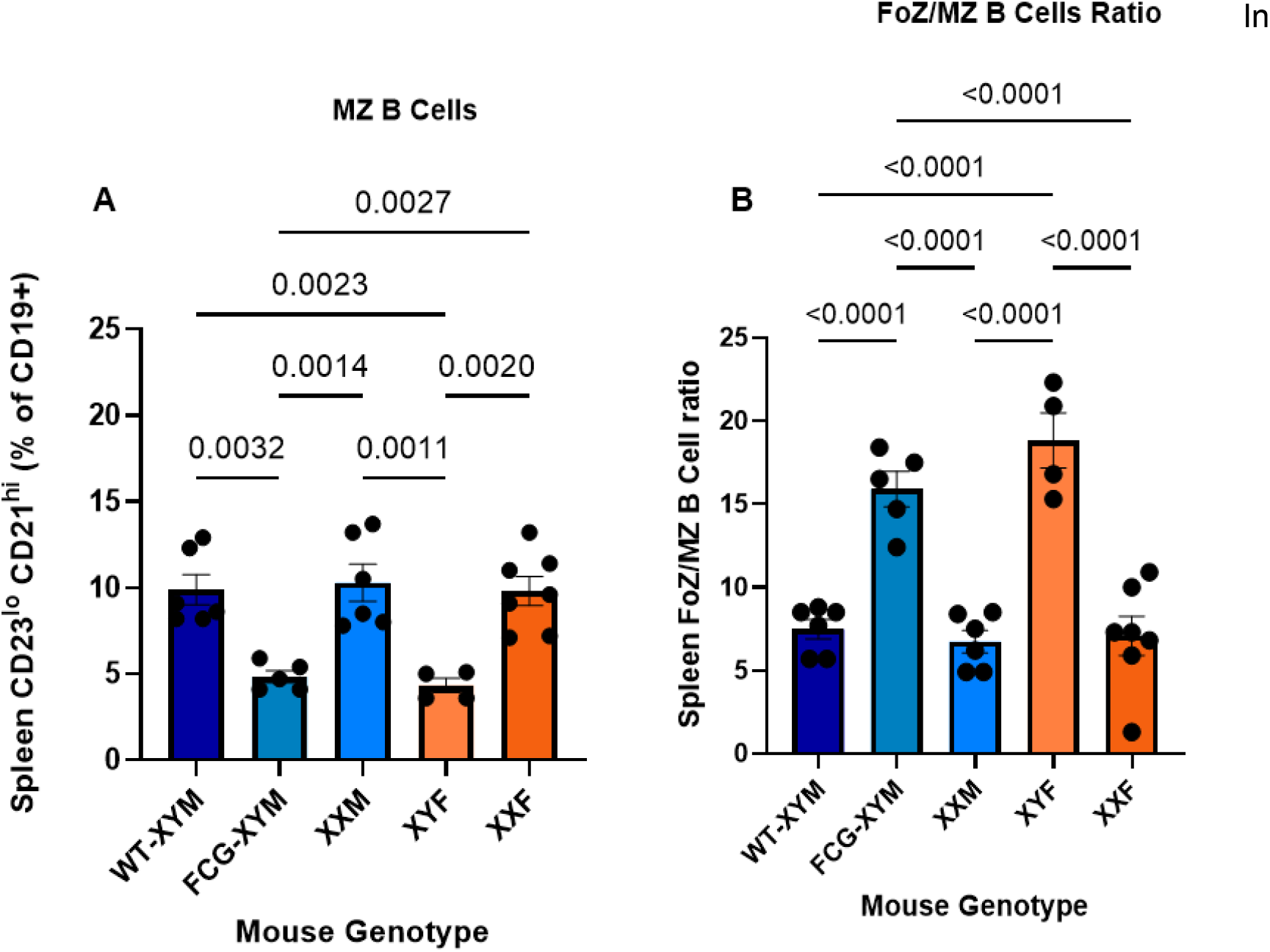
FoB & MZB B cells in spleen between Four Core Genotypes (FCG) mice and wildtype (WT) C57BL/6J males. (A) MZ B cells in spleen (CD23^lo^ CD21^hi^, % of CD19+), and (B) FoZ/MZ B cell ratio in spleen. For (A) and (B): N_WTM_=6, N_XYM_=5, N_XXM_=6, N_XYF_=4, N_XXF_=7. Male gonadal sex marked in blue; female gonadal sex marked in orange. All stats One-way ANOVA plus Tukey-HSD on the means (significance at p<0.05), error bars SEM. Significance bar not shown defines p≥0.05, which is nonsignificant (NS).

### FCG Mice Highlight the Significant Role of Sex Steroid Hormones, and not Host Sex Chromosome Complement, *in vivo* in Adult Peripheral T Cell Subpopulations

The FCG model allows one to determine if the data is influenced by the sex steroid hormones, the sex chromosome complement, or both^39^. So, by stratifying the peripheral T lymphocyte counts within the FCG model, we investigated if there was any gonadal or sex chromosome effects on the mouse that we could not see in earlier analyses just comparing to wildtype (**Figure 9 and 10**).

**Figure 9:**
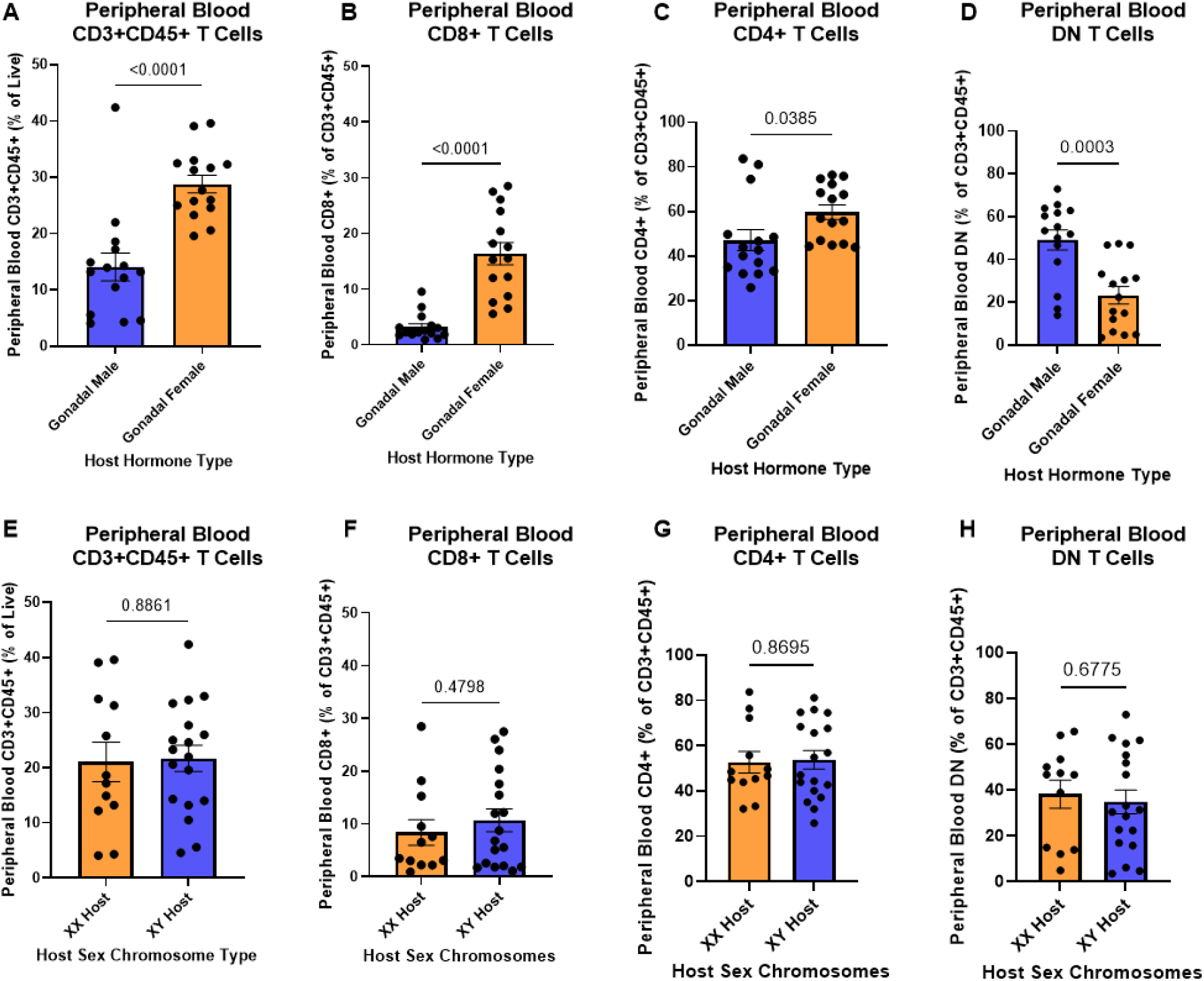
One-way analysis of the effect of gonadal hormones and host sex chromosome complement on peripheral blood T cell subpopulations in FCG mice, in vivo. For (A)-(D), analysis by gonadal sex and hormone axis in the FCG mice. For (E)-(F), analysis by host sex chromosome complement in the FCG mice. (A) and (E) Peripheral blood CD3+CD45+ total T lymphocytes (% of live), (B) and (F) peripheral blood CD8+ T lymphocytes (% of CD3+CD45+), (C) and (G) peripheral blood CD4+ T lymphocytes (% of CD3+CD45+), and (D) and (H) peripheral blood DN T lymphocytes (% of CD3+CD45+). For (A) to (D): N_male_=15, N_female_=15. For (E) to (H): N_XX_=12, N_XY_=18. Male gonadal sex and XY host marked in blue; female gonadal sex and XX host marked in orange. All stats One-way ANOVA plus Tukey-HSD on the means (significance at p<0.05), error bars SEM. Significance bar not shown defines p≥0.05, which is nonsignificant (NS).

**Figure 10:**
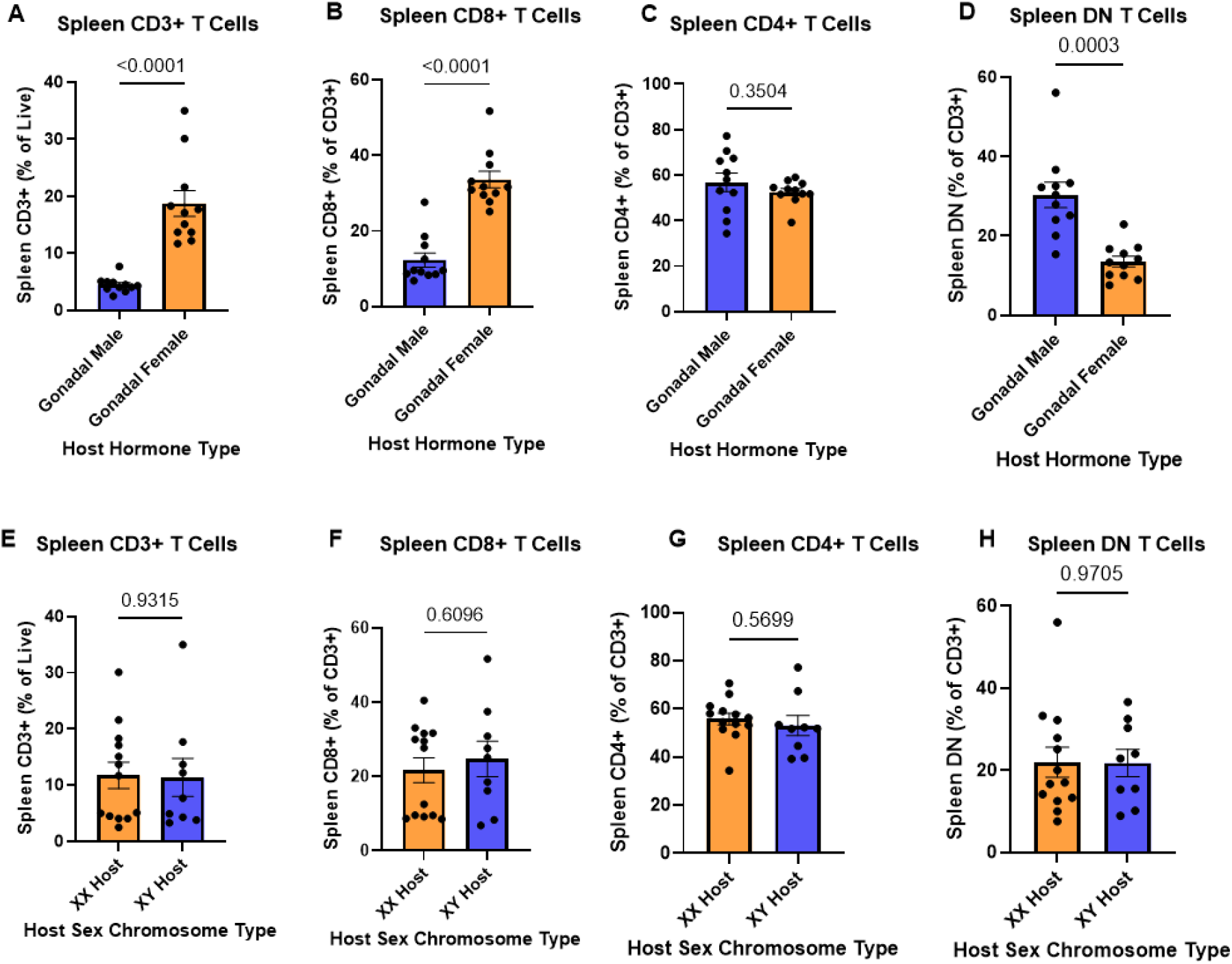
One-way analysis of the effect of gonadal hormones and host sex chromosome complement on spleen T cell subpopulations in FCG mice, *in vivo*. For (A)-(D), analysis by gonadal sex and hormone axis in the FCG mice. For (E)-(F), analysis by host sex chromosome complement in the FCG mice. (A) and (E) spleen CD3+ total T lymphocytes (% of live), (B) and (F) spleen CD8+ T lymphocytes (% of CD3+), (C) and (G) spleen CD4+ T lymphocytes (% of CD3+), and (D) and (H) spleen DN T lymphocytes (% of CD3+). For (A) to (D): N_male_=11, N_female_=11. For (E) to (H): N_XX_=13, N_XY_=9. Male gonadal sex and XY host marked in blue; female gonadal sex and XX host marked in orange. All stats One-way ANOVA plus Tukey-HSD on the means (significance at p<0.05), error bars SEM. Significance bar not shown defines p≥0.05, which is nonsignificant (NS).

In the peripheral blood, there are significant differences between FCG gonadal females and gonadal males (**Figure 9 A-D**). The overall CD3+CD45+ lymphocyte count is higher in gonadal females than in gonadal males by 50% (**Figure 9-A**). This corresponds to an estimated 80% increase in the CD8+ count in gonadal females (**Figure 9-B**). Inversely, we see that gonadal males show a 50% increase in the DN T cell population (**Figure 9-D**). This recapitulates what was seen in the broader analysis of FCG and wildtype XY males and highlights that gonadal females and wildtype XY males have similar T cell counts in peripheral blood. One finding that emerges when looking at differences within the FCG is that there is a slight but significant increase in the CD4+ T cells circulating in peripheral blood in gonadal females (**Figure 9-C**). This was not expected from earlier data. Though there are differences between host hormonal types, we do not see any significant differences based on host sex chromosome complement (**Figure 9 E-H**). Regardless of if a host is XX or XY, there are no significant differences in the T cell count. When looking at the spleen, we see similar results (**Figure 10**). The overall T cell count is increased by 80% in gonadal females (**Figure 10-A**), corresponding to an increase of 60% in the CD8+ T cell population in gonadal females (**Figure 10-B**). Gonadal males have a significant increase in their DN T cell count by 50% (**Figure 10-D**). However, even though the circulating levels of CD4+ T cells seems to be different by gonadal sex, and thus hormone dependent, there are no significant differences in splenic CD4+ T cell counts between gonadal males and females in the FCG model (**Figure 10-C**). All changes are significant between hormone type of the host and not host sex chromosome complement (**Figure 10 E-H**).

## Discussion

The Four Core Genotypes (FCG) mouse model has been widely used to disentangle the effects of sex hormones and sex chromosome complement independently, as well as the interaction of these factors^31,39^. Canonical male versus female comparisons do not allow for this kind of analysis, because sex chromosome effects often co-vary with hormonal effects^39^. Therefore, the FCGs model is being increasingly used to dissect the factors contributing to sex differences. Differences in immunological phenotypes in the FCG model have been previously reported, affecting susceptibility to autoimmune disease^6,43–46^, responses to viral and bacterial pathogens^47–49^, and immune cell subset gene expression, distribution and differentiation^50–52^. The model has also been used in bladder cancer^34^.

Our study reveals that FCG XY and XX males (bearing a multi-copy *Sry* transgene on chromosome 3^39,40^) deviate from wildtype XY males with respect to total and CD8+ peripheral T lymphocytes in the peripheral blood (**Figure 4**), the spleen, and LN (**Figure 5**). In contrast, peripheral T lymphocyte populations in wildtype XY males are similar to both FCG XX and XY females. This suggests an effect of the *Sry* transgenes on T lymphocytes in circulation and in the peripheral lymphoid organs. Our findings are consistent with and expand on those reported specifically in popliteal lymph nodes of wildtype and FCG males^50^, and in gonadectomized FCG males and females^51^.

Previous reports suggested a link to transgenic *Sry* overexpression and/or hormonal regulation of *Glycam1*, a T cell trafficking gene in lymph nodes^50^. We find that there are lower T lymphocytes, specifically in CD8+ cell fractions, which was previously observed in gonadectomized and 7-day-old FCG mice, suggesting a global T cell defect causing these differences^51^. We also observed a minor but statistically significant difference in FCG XY male thymic CD8+ and CD4/CD8 double positive (DP) T cells (**Figure 7**), which also suggests a perturbation of early T cell development, though more comprehensive studies and a time course on younger mice would be needed to confirm this possibility.

The *Sry* transgenes could influence baseline T cell immunity by disrupting the transcription of neighboring genes on chromosome 3, and it is already known that two genes on chromosome 3 are altered in the liver in FCG mice with the *Sry* gene insertion^33^. Some of the genes on chromosome 3 have immune function, for example *Trim59, Il12a, Rarres1, Gfm1*, and *Mlf1*^*33*^. Future studies will be aimed at assessing the expression of *Sry* and of its neighboring genes on chromosome 3, to determine the effects of the transgene on the transcriptome of immune cells.

The reduction of CD8+ T cells in *Sry*-transgenic FCG mice (males) is predicted to have a significant impact on immune responses, including responses to intracellular pathogens and tumors^53^. However, it has been previously reported that susceptibility to pathogens, such as influenza A, have been reported to be essentially the same between FCG males and females, though there was a notable difference in splenic Th1-phenotype cells^49^. In bladder cancer models, FCG males showed increased tumor growth compared to females and similar growth compared to wildtype XY males, concluding that growth differences were dependent on gonadal hormones^54^. Therefore, the previous reports suggest that there may not be a major impact on anti-viral and anti-tumor response in the context of these models, though more functional studies must be done to fully assess the consequences of these differences.

Our analysis of T cell subpopulations also shows that FCG males display a significant increase in the fraction of peripheral double negative (DN) T cells. DN subsets typically constitute only 5% of all peripheral T cells in humans and are about 1-3% in the mouse^55^. Their origin is not completely understood and may originate from escape of negative selection events in the thymus^56^. An expansion of this population was originally considered an abnormal finding, but they have been found to participate in normal physiology^55,56^. *Fas/Fas* ligand deficiency in mice and human (autoimmune lymphoproliferative syndrome) leads to lymphadenopathy and splenomegaly and accumulation of DN cells in peripheral blood and secondary lymphoid organs^56^. DN T cell expansion is also observed in other autoimmune conditions such as Systemic Lupus Erythematosus, Sjogren’s Syndrome, Psoriasis, and in several cancers, i.e. non-small cell lung cancer (NSCLC)^55^. DN cells exhibit broad cytokine expression patterns and show spontaneous release of Il-10 which may be linked to effector and regulatory function^56^. They also play a major role in responses to infection^55^. Interpreting the mechanistic origin and function of this baseline expansion is important in the use and results produced from the FCG model.

The other major differences observed among FCG mice relate to the decrease in marginal zone (MZ) B cells in Y^**-**^ chromosome-bearing male and female mice compared to their XX littermates. The 9 genes known to be translocated in FCG mice from the X chromosome onto the Y chromosome are *Tmsb4x, Tlr8, Tlr7, Prps2, Frmpd4, Msl3, Arhgap6, Amelx*, and *Hccs*^*35*^. While the translocated genes have not been found to be aberrantly overexpressed in all tissue types^38^, and the FCG model’s intrinsic controls can minimize interpretation issues, immune system effects are clearly possible due to overexpression of at least some of the transgenic genes in the spleen of FCG Y-bearing mice^35^. For instance, transgenic overexpression of *Tlr7* alone has been shown to be associated with lower MZ B cell numbers^57^, and may account for the observed phenotype. In addition, the same 9 genes in the FCG Y-translocated region, plus 6 additional ones, characterize the murine Yaa translocation that is associated with lupus development and a reduction to MZ B cells^58–60^. Paradoxically, it has been shown that Y^**-**^ chromosome FCG mice had decreased lupus severity in the SJL and NZM background^6,46^, so the genetic effects underlying these immune phenotypes are likely complex and need more mechanistic and functional follow-up. The impact of this is that MZ B cells play a significant role in responses to T-independent antigens (e.g., bacterial polysaccharides and lipids), such as during responses to *Streptococcus pneumoniae*^*61,62*^, and therefore the effects of the Y^**-**^ translocation on MZB numbers may account for at least some of the sex-chromosome-related differences in responses to heat-killed pneumococci observed in Y^**-**^ chromosome-bearing mice^47,48^.

Our studies add to the characterization of immune phenotypes associated with genetic alterations present in the FCG model background. They highlight the importance and careful use of controls in future studies with this model and help with the interpretation of published results. In addition, further characterization of the effects of these genetic alterations on immune system cells may allow for isolation of new candidate genes and mechanisms involved in immune development, maintenance, and downstream functional consequences. By using internal controls within the FCG model, we identified the major role of the hormone microenvironment, with no detectable effects of the sex chromosome complement in peripheral T cell subpopulations. Additionally, by comparison to wildtype XY males, we see differences due to the *Sry* transgene and in mice that may overexpress 9 genes due to the translocation. Excitingly, this translocation may allow for easier isolation of candidate genes and mechanisms in adult immune maintenance and development.

The FCG model allows for intrinsic control of two independent variables of sex, and when compared to wildtype, allows for detection of differences along the transgene or translocation axis^39^. Future studies detailing the transcriptional and functional consequences of the discrepancies between wildtype and FCG mice will allow the FCGs model to continue to inform studies of critical sex differences in physiologic and pathogenic states.

## Acknowledgements

We acknowledge the work done on peripheral blood flow cytometry and phenotyping by the core directed by Dr. Kerry Campbell at the Fox Chase Cancer Center.

## Availability of Data and Materials

Raw data and FSC files from flow cytometry will be available upon request and will be attached as supplementary when published in a peer-reviewed journal. Future sequencing will be also available upon request and available publicly on the NIH-NCBI Gene Expression Omnibus (GEO) for download.

## Declarations

Authors have no declarations or conflict of interests to disclose

